## Additional file 1 for "BindVAE: Dirichlet variational autoencoders for *de novo* motif discovery from accessible chromatin"

Meghana Kshirsagar*
Microsoft, AI for Good Research Lab, Redmond, WA, USA.

Han Yuan
Calico Life Sciences, South San Francisco, CA, USA

Juan Lavista Ferres
Microsoft, AI for Good Research Lab, Redmond, WA, USA.

Christina Leslie*
Memorial Sloan Kettering Cancer Center, New York, NY, USA

June 18, 2022

#### 1 Datasets

We show the datasets used in further experiments in Supplemental Table S1.

##### 1.1 CAP-SELEX

Combinations of TFs with successful CAP-SELEX experiment are limited after filtering for quality metrics as described in our BindSpace paper (Nature Methods 2019). Even after filtering for QC, in only a handful ( 70 of them) is there a composite motif that differs from the concatenation of individual motifs for which we can assess enrichment in the latent dimensions. Rather than make systematic statements about composite motifs given the sparsity of CAP-SELEX data, we are only presenting "analysis vignettes" that make the point that in some cases the latent dimension can capture a composite motif.

| Experiment | T cell female adult |
|---|---|
| URL: `https://www.encodeproject.org/experiments/ENCSR977LVI/` | |
| replicate 1, number of peaks | 79802 |
| replicate 2, number of peaks | 82359 |
| overlap in peaks | 68682 |

| Experiment | naive B cell |
|---|---|
| female donor, number of peaks | 87500 |
| URL: `https://www.encodeproject.org/experiments/ENCSR685OFR/` | |
| male donor, number of peaks | 94007 |
| URL: `https://www.encodeproject.org/experiments/ENCSR903WVU/` | |
| overlap in peaks | 76442 |

| Experiment | naive CD8+ T cells |
|---|---|
| mouse, number of peaks | 221,053 |
| URL: `https://pubmed.ncbi.nlm.nih.gov/33891860/` | |
| human, number of peaks | 190,816 |
| URL: `https://pubmed.ncbi.nlm.nih.gov/33891860/` | |

**Table S1:** Details of the ATAC-seq datasets used in the analyses shown in supplementary material (for the experiments shown in Figure S8). Overlap of at least 50bp is considered for computing the overlap rows in the table above.

| | | | | |
|---|---|---|---|---|
| **GGGCGG | CCCCG**C | G**CCCGC | **CCGCCG | **GCCCGC |
| CCGCC**C | CGCCC**C | GC**GCGC | CGCGC**C | C**GCCGC |
| **CCGCCC | CC**CGCC | CGC**CCG | CCGC**CG | CGC**CCC |
| **GGCGGG | C**GCCCC | GCGG**CC | **GCGCGC | **GCGCGG |
| **CCCGCC | **GGCGGC | CCG**GCC | *GGCGGGG | CGCG**GC |
| CCGC**CC | **GCCGCC | CGCC**GC | *CCCCGCC | **GCCGGG |
| C**CGCCC | **CGGGGC | CCCC**CC | CCGCC*CC | **CCCGGC |
| CCC**CCC | **GCCCCG | CGC**CGC | **CCGCGC | CCCCGC*C |
| CCCGC**C | GCCC**CC | GC**CGGC | GCGC**CC | CCCGC*CC |
| GC**CGCC | G**GCCGC | CCG**CCC | C**GGGCG | GC**GGCC |
| C**CCGCC | CC**CCCC | *GGGCGGG | CC*CGCCC | CG**CCGC |
| G**CCGCC | G**GCGGC | *CCCGCCC | C**CGCGC | CCC*GCCC |
| **GCGGGG | C**CCCGC | CC**CCCG | C**CGCCG | **GCCGCG |
| **CCCCGC | G**GGGGC | C*CCGCCC | **GCGGCC | G**CGCGC |
| **GGGGCG | C**CCCCG | CG**GGCG | *GGGGCGG | C*CGCCCC |
| **CGCCCC | GCC**CGC | CGCC**CC | *CCGCCCC | C**GCGGC |
| CC**GCCC | GCG**GCC | CC**CCGC | GC*CCGCC | CCC**CGC |
| CCCG**CC | C**GGCGG | CG**GCCG | CC**GCGC | CCGGC**C |
| GCC**GCC | **CGCCGC | CCCGCC*C | C**GCGCC | **GGCGCG |
| GC**CCGC | **GCGGCG | | CCGC**GC | |

**Fig. S1:** The top 100 8-mers from the latent dimension #37, that captures genomic background in the GM12878 model.

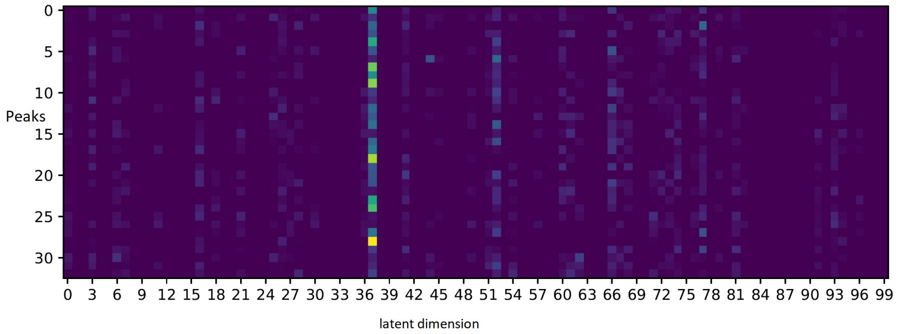


**Fig. S2:** The latent representation learned for top low complexity regions (LCRs) or regions with repeats are similar as shown in the heatmap, where most of them have a high value for the same latent factor (#37).

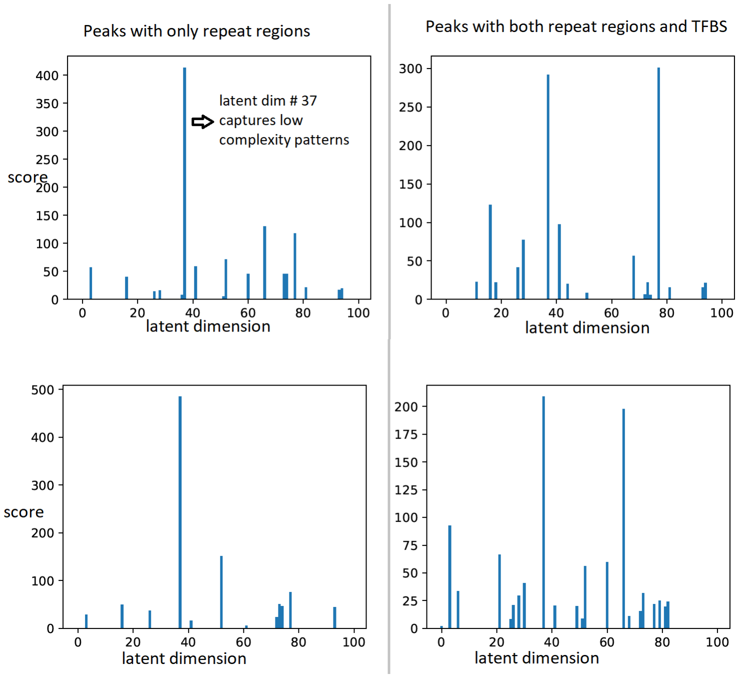


**Fig. S3:** Latent projection of peaks with only repeat regions and peaks with some low-complexity repeating patterns and TFBS for a TF. This shows the disentanglement done by BindVAE.

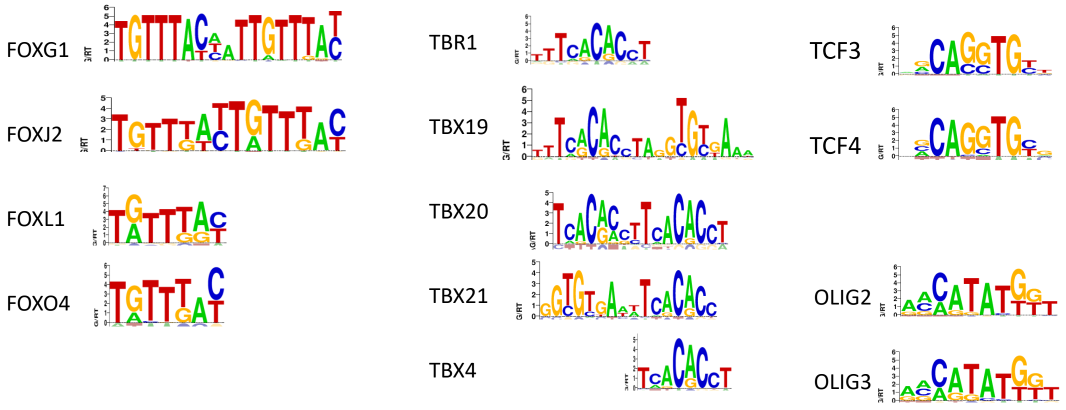


**Fig. S4:** CIS-BP motifs for TFs from the same family or for paralogous TFs are shown, to illustrate the difficulty of learning TF-specific patterns for these. We show the TFs from the heatmap of **Figure 2b** (TFs in the boxes). Each group of TFs gets projected to the same latent factor by our model as discussed in the main paper.

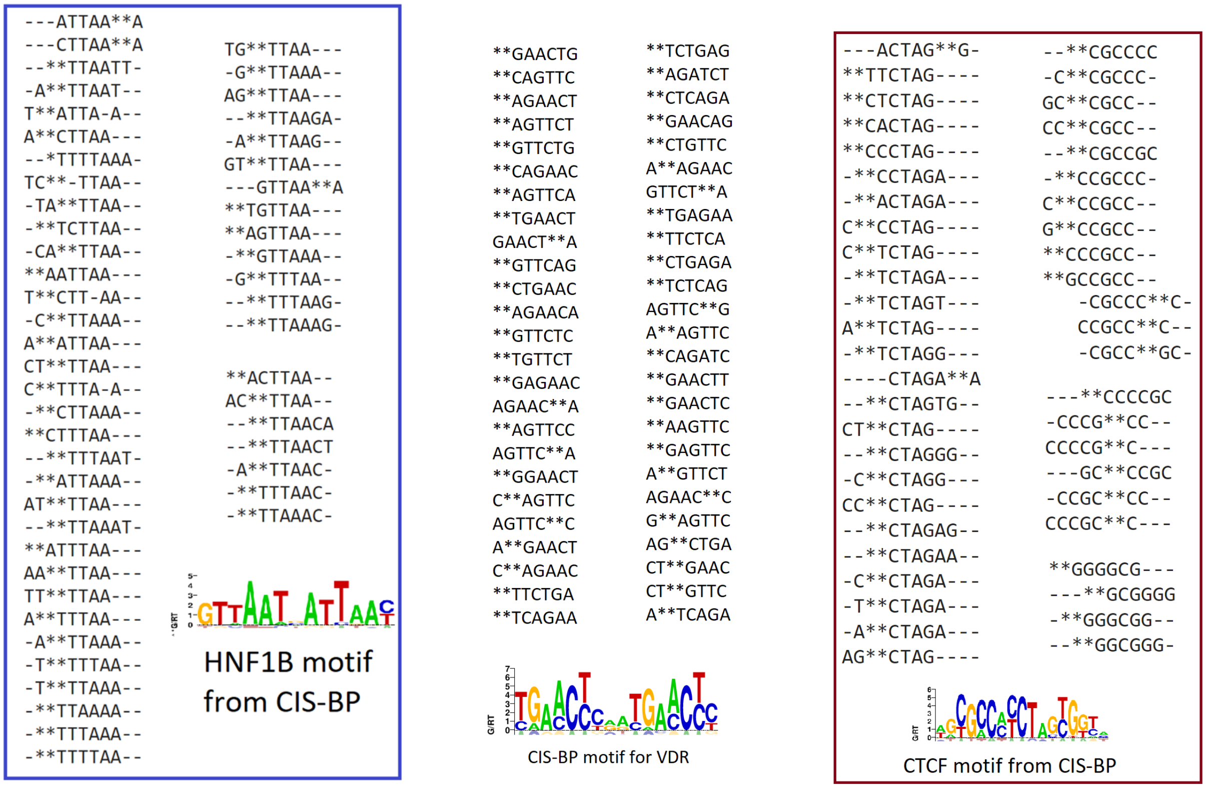

**Fig. S5:** GM12878: Top 50 8-mers from some latent dimensions, aligned using clustal omega to summarize the patterns found. The CIS-BP motif corresponding to the TF that was assigned to each latent dimension (using Algorithm 1) is also shown. Since CTCF is assigned to multiple latent dimensions, the top 25 8-mers from each are shown.

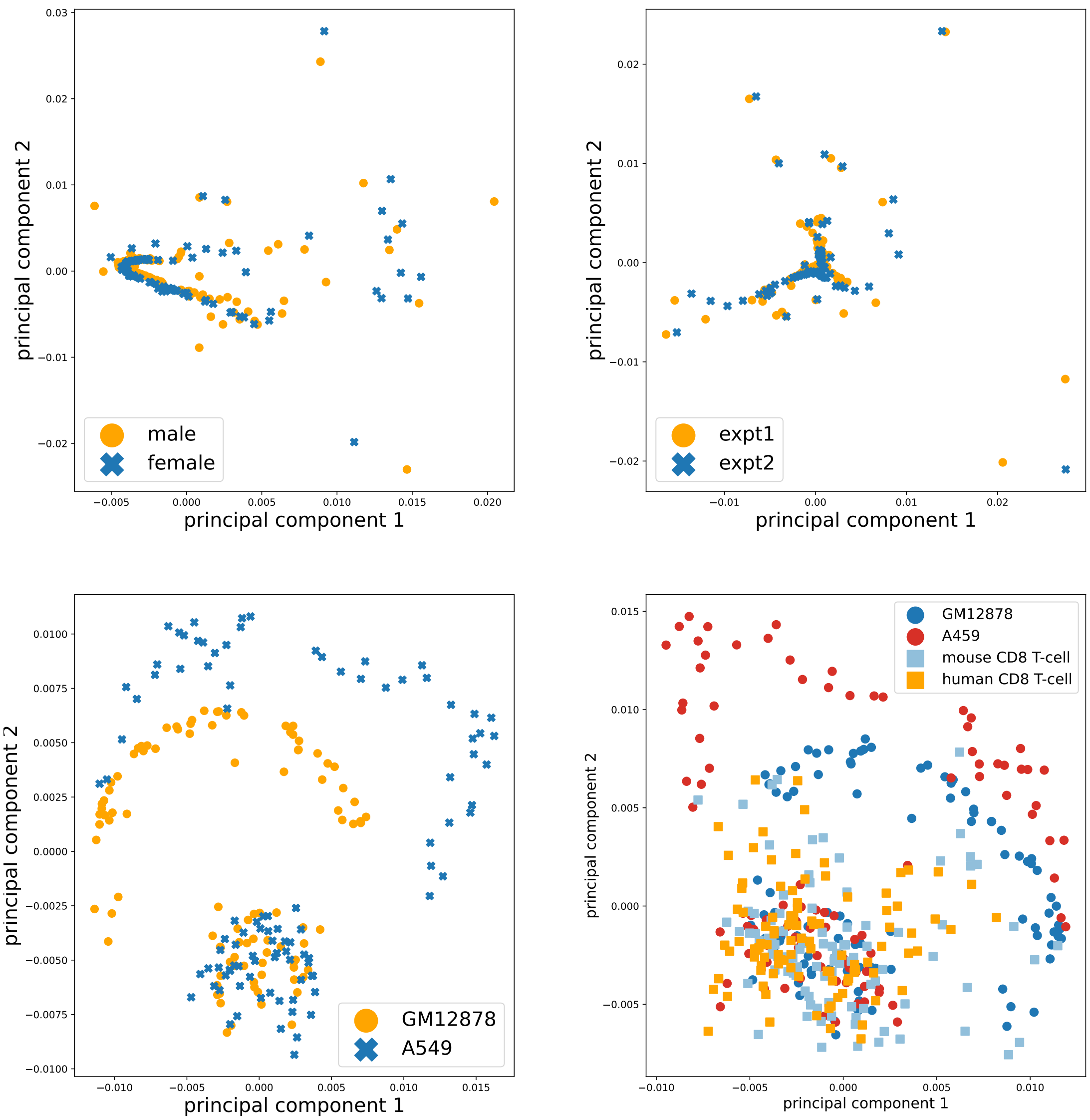


**Fig. S6:** PCA done on the decoder parameters $\theta \in \mathbb{R}^{M \times D}$, that capture the k-mer distributions, from models trained on various cell-types. Each dot on a plot represents a TF binding pattern or k-mer distribution. **(Top left)** k-mer distributions from two models trained on naive B cells from two human donors, male and female. **(Top right)** k-mer distributions from two models from two repeat experiments on female naive B cells. In both cases, there is not much variance in the learned patterns, as these are biologically close samples. **(Bottom left)** k-mer distributions from the two models trained on GM12878 and A549. Since these are distinct cell types, we see two distinct manifolds in the patterns being learned. There is also a cluster of latent dimensions at the bottom of the plot that captures similar k-mer patterns. **(Bottom right)** k-mer distributions from four models: GM12878, A549, mouse CD8 T cells and human CD8 T cells.

| TF | GM12878 motif | A549 motif | CIS-BP motif |
|---|---|---|---|
| HNF4A | 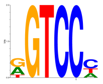 | 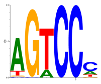 | 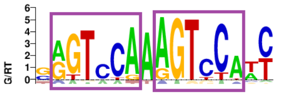 |
| NFIA | 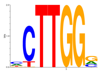 | 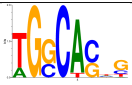 | 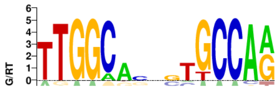 |
| SRY | 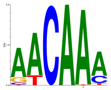 | 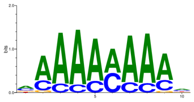 | 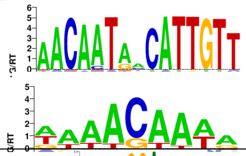 |
| ELF5 | 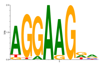 | 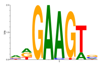 | 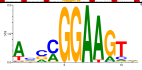 |
| OLIG3 | 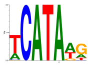 | 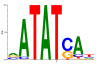 | 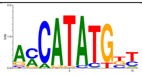 |

**Table S2:** Motifs constructed from latent factors learned for GM12878 and A549 for TFs learned by our model. `HNF4A`, `NFIA`, `SRY` have the same 'accessibility score' in both GM12878 and A549 trained models (see **Figure 3d** in the main paper). `ELF5` and `OLIG3` are both expressed in both cell types.

### 2 Comparison to baselines

**MEME:** We ran MEME in tandem with Tomtom, where the learned motifs from MEME are used as input to Tomtom which matches the learned motifs with the PWMs from our dataset of 270 HT-SELEX TFs. We used the default parameters to run MEME, except for the following: number of motifs=500, background-model=2nd order, motif distribution=anr, no e-value threshold. The following command was used:

```
./meme -dna -mod anr -nmotifs 500 -minw 5 -maxw 15 -markov_order 2 -p
10 <fasta-file>
```

MEME was run with 10 processors in parallel. On both datasets, MEME+Tomtom took 10 to 12 hours to run, with Tomtom taking under 5 minutes.

**GADEM:** Overall, this genetic algorithm guided approach is non-deterministic (produces differing number of motifs in each run) and suited for learning a single TF's motif/PWM from ChIP-seq data. Since GADEM can only find motifs, we ran it in conjunction with Tomtom for motif-matching to find which SELEX PWMs are found. The R-package rGADEM had some limitations (can only work with 44000 sequences, limited number of motif matches etc.); we modified the source code as per the documentation and ran it on the GM12878 and A549 datasets. We used the default

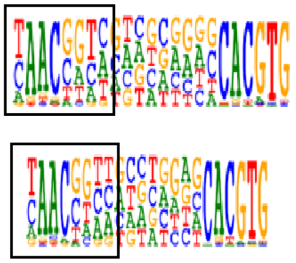

Motif from Jolma et al.

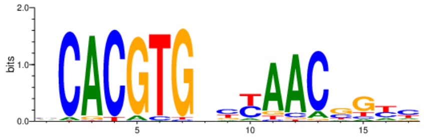


Motif generated by MEME using enriched CAP-SELEX probes

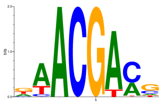

BindVAE motif

| | Other cooperative TF pairs found | |
|---|---|---|
| ELK1_SPDEF | ELK1_TCF15 | MYBL1_MAX |
| ELK1_NFKB1 | TFAP4_FLI1 | TFAP4_MAX |
| E2F1_NHLH1 | ETV2_NHLH1 | HOXB2_NHLH1 |
| POU2F1_FOXO6 | E2F1_ELK1 | MYBL1_MAX |
| FOXJ3_TBX21 | | |

**Table S3:** MYBL1:MAX motifs computed from the GM12878 model are shown in the top row. The bottom table shows all cooperative binding pairs of TFs found for GM12878 by our model using the CAP-SELEX data are shown.

| Method | Run-time | GM12878 | | | A549 | | | T cells female adult | | |
|---|---|---|---|---|---|---|---|---|---|---|
| | | Prec. | Rec. | F1 | Prec. | Rec. | F1 | Prec. | Rec. | F1 |
| GADEM+Tomtom | 2-7 hrs | 0.634 | 0.154 | 0.247 | 0.571 | 0.090 | 0.154 | 0.760 | 0.186 | 0.298 |
| MEME+Tomtom | 9-10 hrs | 0.530 | 0.261 | 0.348 | 0.791 | 0.327 | 0.462 | 0.686 | 0.361 | 0.472 |
| HOMER | 6-24 hrs | 0.608 | 0.351 | 0.444 | 0.765 | 0.504 | 0.607 | 0.659 | 0.811 | 0.728 |
| BindVAE | 4-5 hrs | 0.636 | 0.416 | 0.502 | 0.806 | 0.359 | 0.496 | 0.785 | 0.351 | 0.485 |
| BindVAE+Tomtom | 4-5 hrs | 0.573 | 0.952 | 0.714 | 0.733 | 0.900 | 0.807 | 0.638 | 0.821 | 0.718 |

**(a)**

| Method | p-value / e-value | GM12878 | | | A549 | | | T cells female adult | | |
|---|---|---|---|---|---|---|---|---|---|---|
| | | P | R | F1 | P | R | F1 | P | R | F1 |
| All positive classifier | | 0.562 | 1.000 | 0.720 | 0.736 | 1.000 | 0.828 | 0.628 | 1.000 | 0.771 |
| HOMER | 0.001 | 0.631 | 0.386 | 0.478 | 0.783 | 0.377 | 0.508 | 0.780 | 0.414 | 0.540 |
| HOMER | 0.01 | 0.608 | 0.351 | 0.444 | 0.765 | 0.504 | 0.607 | 0.659 | 0.811 | 0.728 |
| HOMER | 0.1 | 0.628 | 0.452 | 0.525 | 0.777 | 0.350 | 0.481 | 0.748 | 0.505 | 0.602 |
| HOMER | 10 | 0.719 | 0.458 | 0.559 | 0.800 | 0.418 | 0.548 | 0.764 | 0.430 | 0.550 |
| BindVAE | 3%, 0.005 | 0.744 | 0.208 | 0.324 | 0.833 | 0.113 | 0.198 | 0.821 | 0.164 | 0.273 |
| BindVAE | 5%, 0.05 | 0.636 | 0.416 | 0.502 | 0.806 | 0.359 | 0.496 | 0.785 | 0.351 | 0.485 |
| BindVAE | 5%, 0.1 | 0.636 | 0.541 | 0.584 | 0.779 | 0.513 | 0.618 | 0.766 | 0.436 | 0.554 |
| BindVAE | 10%, 0.5 | 0.579 | 0.803 | 0.672 | 0.747 | 0.754 | 0.750 | 0.721 | 0.611 | 0.661 |
| BindVAE | 10%, 0.8 | 0.561 | 1.000 | 0.718 | 0.735 | 1.000 | 0.847 | 0.765 | 0.382 | 0.509 |
| BindVAE+Tomtom | 0.1 | 0.533 | 0.047 | 0.086 | 0.750 | 0.040 | 0.075 | 0.541 | 0.038 | 0.07 |
| BindVAE+Tomtom | 1 | 0.539 | 0.410 | 0.464 | 0.737 | 0.204 | 0.318 | 0.602 | 0.481 | 0.534 |
| BindVAE+Tomtom | 10 | 0.573 | 0.952 | 0.714 | 0.733 | 0.900 | 0.807 | 0.633 | 0.796 | 0.705 |
| BindVAE+Tomtom | 20 | 0.571 | 0.976 | 0.720 | 0.731 | 0.954 | 0.827 | 0.638 | 0.821 | 0.718 |

**(b)**

**Table S4:** Precision, recall, F1 achieved by the various *de novo* motif discovery approaches in retrieving TFs from ATAC-seq peaks of the two cell types. Expressed TFs (from RNA-seq data) that intersect with our HT-SELEX set of TFs are used as the gold-standard for retrieval. **(a)** Performance with default p-value and e-value cut-offs for all methods. **(b)** For the two best approaches, HOMER and BindVAE, performance upon varying the cut-offs is shown. The performance of a naive match-every-motif classifier (label everything positive) is shown.

parameters (e-value threshold=0), except for the number of motifs to find, which we set as: nmotifs=500. We tried several p-value cut-offs, including the recommended one (0.0002) and less stringent cut-offs. Relaxing the p-value cut-off to values higher than 0.001 results in millions of motif matches being found and the code segmentation faults. Oddly, fewer motifs were found for the less stringent cut-offs of 0.001, 0.0008. The algorithm had run times between 2 to 7 hours (with lower times for more stringent p-value cut-offs). With a cut-off of 0.0002, 4 to 8 motifs were found ranging in length from 9 to 31.

**BindVAE + Tomtom**: In order to understand the contribution of the VAE and Algorithm 1 to the performance separately, we replace Algorithm-1 with a PWM matching approach like Tomtom that takes PWMs generated for each latent dimension and matches them against a database. This gives us another version of BindVAE.

In general, we find that BindVAE has a higher precision compared to other approaches. Replacing the TF-matching procedure (from Algorithm 1) with Tomtom, we find results in a higher recall and F1-score. HOMER has a better performance on the T cell dataset.

#### 2.1 Varying the cut-off thresholds

All the motif finding algorithms and our TF-matching algorithm (Algorithm 1) assign p-values to the matches, which are not appropriate for use as 'scores' from a classifier for generating ROC curves. We show results for the two best performing models: BindVAE and HOMER, for a few different p-value / e-value cut-offs and show the results in **Table S4(b)**.
**HOMER:** The way to achieve more comparisons with HOMER is by changing the e-value cut-off (the default value is: 0.01). This requires re-running the entire algorithm, which is also the case for other motif discovery approaches.
**BindVAE:** being a machine-learning model only requires to be trained once. The p-value changes need to be made in the post-processing in Algorithm 1, which takes 5 mins to complete. There are two thresholds that can be changed:

- the p-value cut-off on line-8 of Algorithm 1 or
- the top% of all probes where we look for enrichment of a TF's probes on line-7

**BindVAE + Tomtom**: Here also, two thresholds can be changed: the e-value cut-off of Tomtom or the parameters of the MEME algorithm, which we use to generate PWMs from 10mers. We do not change the MEME parameters however and only modify the e-value threshold of Tomtom.
**All positives classifier:** The performance of a naive match-every-motif classifier (label everything positive) is shown. This gives a sense of the class skew.

Overall, we find that BindVAE+Tomtom has a higher recall and F1-score compared to HOMER (if one were to plot the area under PR curve, it would be expected to be higher as well).

### 3 Latent factors capturing non-TF related patterns

**Noisy and redundant dimensions in the model:** The model learned on GM12878 peaks has 10 dimensions that capture 'noise' and do not contain any TF-specific information. The k-mer distributions for the noisy latent factors are mostly flat (almost uniform distributions). When DNA sequences are projected onto these dimensions, the resultant matrix has small and constant-valued columns. See Figure S11, where we show the ATAC-seq peaks from our GM12878 dataset projected onto the 10 noisy dimensions from this model.

Redundant dimensions are ones that capture kmer patterns for the same TF. In Figure S10, we show the kmer distributions of two latent dimensions that were mapped to

`HEY1`. Similar distributions are shown for `NFIA` and `TFAP4`. This redundancy is likely due to binding sites of homologous TFs that are not part of our HT-SELEX dataset (which is used by Algorithm-1 to map latent dimensions to TFs).

**Low-complexity regions and genomic background:** We use Tandem Repeat Finder (TRF) to find GM12878 peaks that contain low-complexity regions (LCRs): i.e. regions with an abundance of short tandem repeats or regions with an abundance of a single base (like `AAAAAAAAAA`). See Figure S12 for example sequences found in the GM12878 ATAC-seq peaks. We find that the top scoring 30 peaks (as per scores assigned by TRF) contain repeats throughout the 200bp region, while the subsequent peaks contain some TFBS in addition to some repeat regions. We find that the latent representation learned for these top LCRs or regions with repeats are similar as shown by the heatmap below, where most of them have a high value for the same latent factor (#37). We show the top 100 k-mers from latent factor #37 in Figure S1 and they contain recurring CG-rich k-mers.

To depict the learned theta matrix (k-mer distribution matrix) that is trained on genomic background regions, we trained two models:

1. (Model 1) Random regions of naked DNA from mouse germ cells. We use the following dataset to train this model: `https://www.ncbi.nlm.nih.gov/geo/query/acc.cgi?acc=GSM1550786`

2. (Model 2) flanking regions of GM12878 peaks that are 10kb away

As shown in Figure S13, we find that only a couple of distinct k-mer patterns are learned by Model 1. On the other hand, Model 2 does learn several distinct TF-like signatures similar to the GM12878 model trained on peaks only. We believe this to be the case because distal regions of “promoter peaks” also contain binding sites, and our flanking set of peaks is likely to contain some of these.

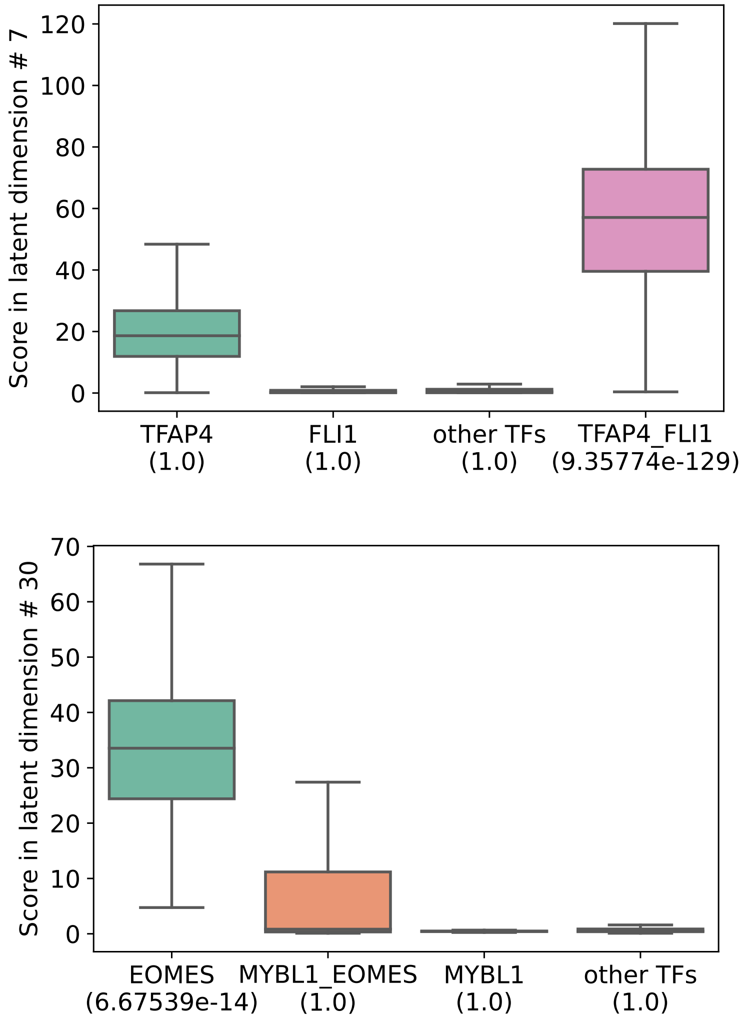


**Fig. S7:** Examples of CAP-SELEX probes scoring higher and lower than individual TF probes. **(Top)** Example of cooperative binding CAP-SELEX probes of TFAP4:FLI1 being enriched while the individual TF probes are not enriched for TFAP4 or FLI1 ($p = 1$). **(Bottom)** Example where individual TF probes from the SELEX experiment for EOMES are enriched, while CAP-SELEX probes of cooperative binding between MYBL1:EOMES are not.

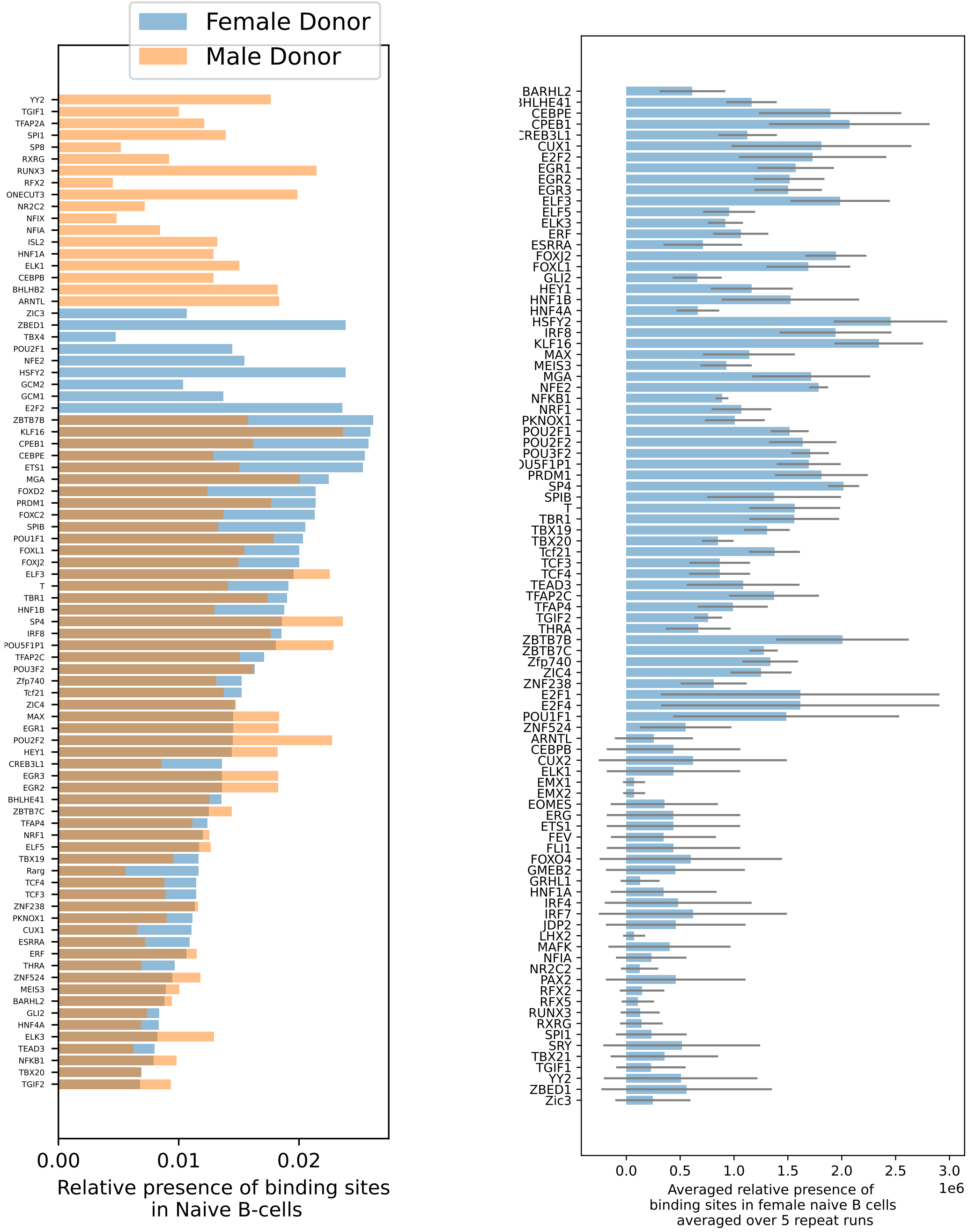


**Fig. S8:** Accessibility scores of TFs obtained by summing the latent representations over all ATAC-seq peaks showing the possible extent of accessibility for each TF. The datasets used here are described in Table S1. **(Left)** TFs found in naive B cells by our model and their ‘accessibility scores’ in the two human donors: male donor (light blue) and female donor (orange).**(Right)** Averaged relative accessibility scores in female donor over 5 runs.

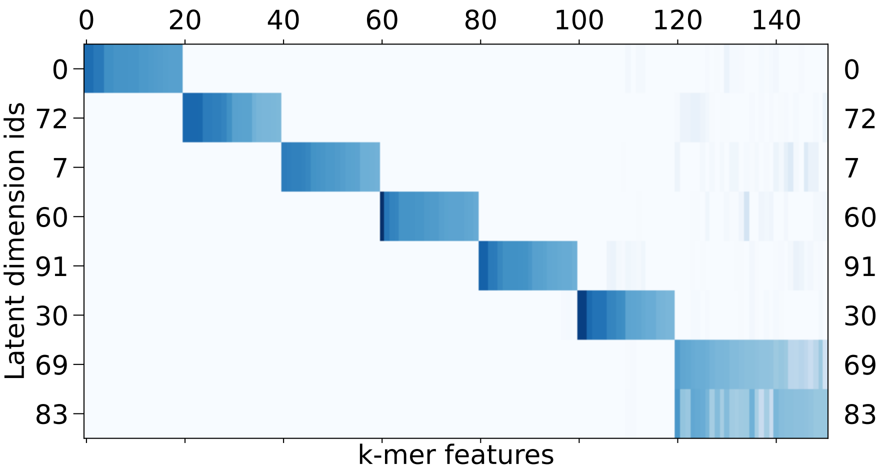


**Fig. S9:** Select latent dimensions (topics) from the GM12878 model, that are referred to in the main text. The shown matrix is a sub-matrix of $\theta \in \mathbb{R}^{M \times D}$. Dimensions 69 and 83 are redundant, in that they both assign high weights to the same k-mer features.

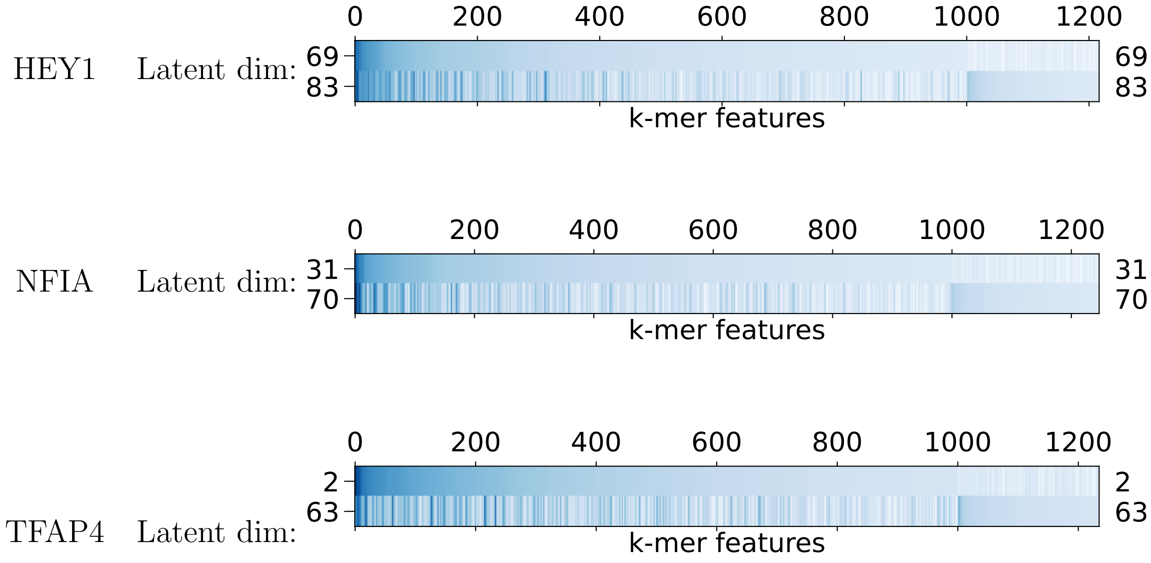


**Fig. S10:** Redundant dimensions: Latent dimensions with similar k-mer distributions are shown. In each plot, the two dimensions were mapped to the same TF; for example: #69 and #83 are both mapped to HEY1. Along the x-axis is the union of the top 1000 8-mers from both dimensions. The values in each row are the decoder parameters learned by the model: $\vec{\theta}_i$ for the $i^{th}$ dimension.

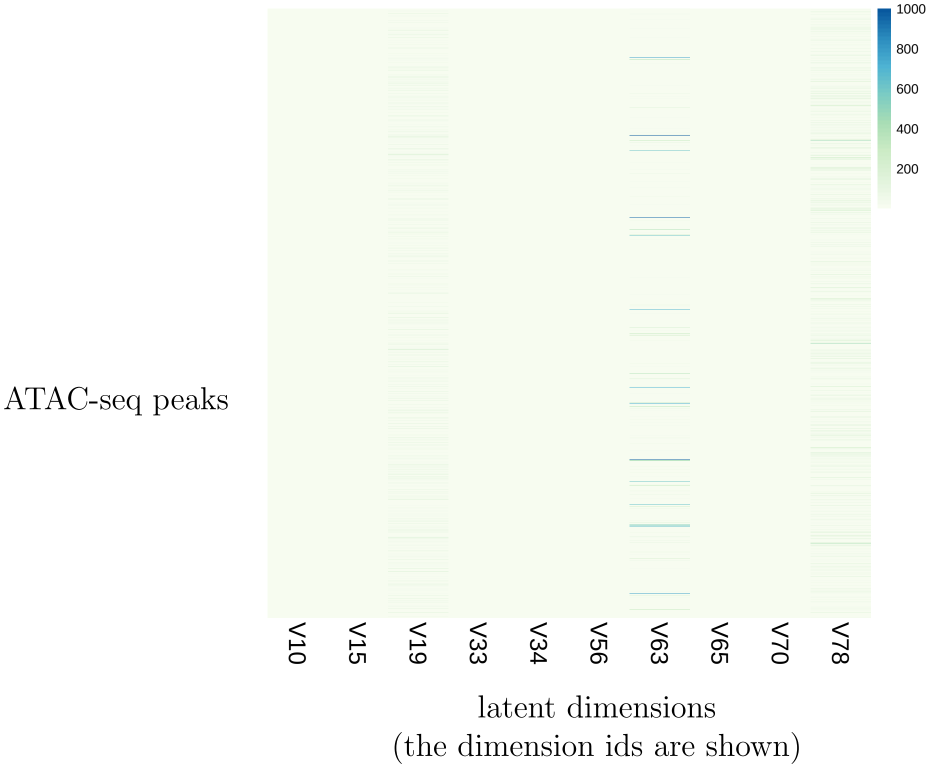


**Fig. S11:** GM12878 peaks projected onto the 10 noisy latent dimensions.

| Learned TFs that are expressed | Not expressed |
|---|---|
| CPEB1 | ALX1 |
| CREB3L1 | AR |
| CTCF | BHLHE23 |
| CUX1 | BHLHE41 |
| CUX2 | EGR4 |
| E2F2 | ELF5 |
| EGR3 | EMX2 |
| ELK1 | ERG |
| ELK3 | ESRRG |
| EMX1 | FEV |
| EOMES | FOXC2 |
| ERF | FOXL1 |
| ESRRA | GATA4 |
| ETS1 | GATA5 |
| FOXD2 | GLI2 |
| FOXG1 | HNF1A |
| FOXJ2 | HNF1B |
| FOXO4 | HNF4A |
| GMEB2 | HSFY2 |
| GRHL1 | MEIS3 |
| HMX3 | MEOX2 |
| HSF1 | NFE2 |
| HSF2 | NKX3-2 |
| HSF4 | OLIG2 |
| IRF4 | OLIG3 |
| IRF7 | ONECUT3 |
| IRF8 | PAX7 |
| KLF16 | PKNOX2 |
| LHX2 | POU1F1 |
| MAFK | POU3F2 |
| MAX | POU3F4 |
| MGA | POU5F1P1 |
| MLX | RXRG |
| MLXIPL | SP8 |
| NFIA | SRY |
| NFIB | T |
| NFIX | TBR1 |
| NFKB1 | TBX20 |
| NR2C2 | TBX4 |
| NR2F1 | TCF21 |
| NR2F6 | TCFAP2A |
| NR3C1 | TFAP2C |
| NRF1 | UNCX |
| PKNOX1 | ZBTB7C |
| POU2F2 | ZFP740 |
| PRDM1 | ZIC3 |
| RARG | ZIC4 |
| RFX2 | ZNF238 |
| RFX3 | ZSCAN4 |
| RFX5 | |
| RUNX2 | |
| RUNX3 | |
| RXRA | |
| RXRB | |
| SMAD3 | |
| SPI1 | |
| SPIB | |
| SRF | |
| TBX19 | |
| TBX21 | |
| TCF3 | |
| TCF4 | |
| TEAD3 | |
| TEF | |
| TFAP2A | |
| TFAP4 | |
| TGIF1 | |
| TGIF2 | |
| THRA | |
| VDR | |
| YY2 | |
| ZBTB7B | |
| ZNF524 | |

**Table S5:** TFs learned by the model trained on the GM12878 ATAC-seq data.

chr5:   11488-11687
CTAACCCTAACCCTAACCCTAACCCTAACCCTAACCCTAACCCTAACCCTAACCCTAACCCTAACCCTAACCCTAACC
CTAACCCTAACCCTAAACCCTAACCCTAACCCTAACCCTAACCCTAACCCTAACCCTAACCCTAACCCTAACCCTAAC
CCTAACCCTAACCCTAACCCTAACCCTAACCCTAACCCTAACCC

chr7:   158996234-158996433
CCAGGATGCCCTGTCCATGAGGAGACGGCCTGGGACCAGGATGCCCTATCCATGAGGAGACGGCCTGGGACCAG
GATGCCCTATCCATGAGGAGACAGCCTGGGACCAGGATGCCCTATCCATGAGGAGACGGCCTGGGACCAGGGTG
CCCTATCCATGAGGAGACGGCCTGGGACCAGGGTGCCCTTTCCATGAGGAGA

chr6:   170709160-170709359
AGAACTATGTCATAAGCCTTAGGAGTGAGTGAGAAAGGTAGAACTATGTCATAACCCTTAGGAGTGAGTGGGGA
AGGTAGAACTATGTCATAACCCTTAGGAGTGAGTGAGGAAGGTAGAACTATGTCATAACCCTTAGGAGTGAGTG
GGGAAGGTAGAACTATGTCATAACTCTTAGGAGTGAGTGGGGAAGGTAGAAC

chr18:   10005-10204
AACCCTAACCCTAACCCTTACCCCTAACCCTAACCCTAACCCTAACCCTAACCCTAACCCTAACCCTAACCCTACCCT
AACCCTAACCCTACCCTAACCCTAACCCTAACCCTAACCCTAACCCTAACCCTAACCCTAACCCTTAACCCTAACCCTT
AACCCTAACCCTAACCCTAACCCTAACCCTAACCCTAACCCTA

chr18:   72485313-72485512
GAAGCCAAGTTACTAGAGAGCTTAAAGAAGAAATTGACAATAGTATCAAAGAAGAAGTGATCAGAAGTGGTTTG
GAACTATCAGTAGAAGCCAAGTTATTAGAGAGCTTAAAGAAGAGATTGACAATAGTATGAAAGAGGAAGTGATC
AAAAGTGGTTTGGAACTATCAGTGGAAGCCAAGTTACTAGAGAGCTTAAAGA

chr13:   113527422-113527621
TGTGGCTTTGTCTTCCTGAACGGTGTGAGGTGTGGCTTTGTCTTCCTGAGTGGTGTGAGGTGTGGCTTTGTCTTCC
TGAGCGGTGTGAGGTGTGGCTTTGTCTTCCTGAGTGGTGTGAGGTGTGGCTTTGTCTCCTTAATGGTGTGAGGTG
TGGCTTTGTCTTCCTGAGCGGTGTGAGGTGTGGCTTTGTCTTCCTGAAC

**Fig. S12:** GM12878 peaks that contain sequences with the most repeats (top 5 are shown), as ranked by the Tandem Repeat Finder score (TRF algorithm ). We find that the highest-scoring topic for all of these sequences is the same. The genomic coordinates correspond to the `hg19` assembly.

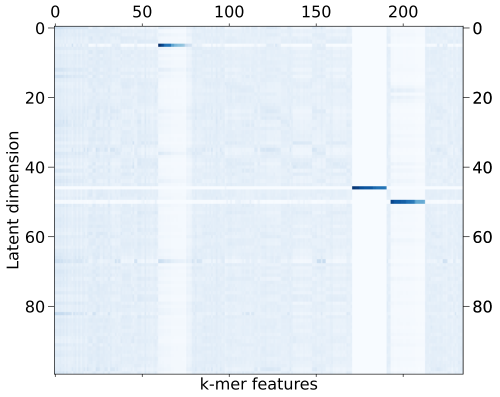


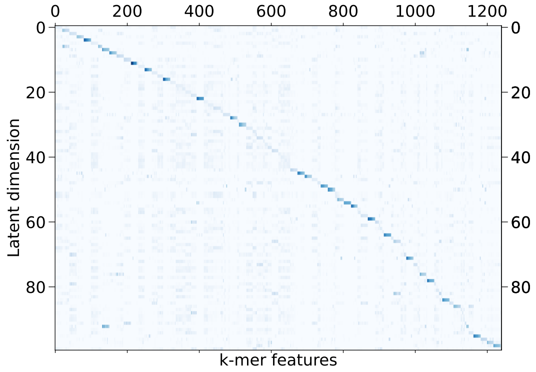


**Fig. S13:** Interpreting the models learned on random regions of the DNA. **(Top)** Matrix showing the learned decoder parameter $\theta$ (that shows the k-mer distributions for each latent factor) for a model trained on 150,000 random regions of naked DNA from mouse E16.5 sorted germ cells. **(Bottom)** Matrix showing the learned decoder parameter $\theta$ for a model trained on flanking genomic regions from GM12878 cells, where regions that are 10kb away from the ATAC-seq peaks were chosen as training data.

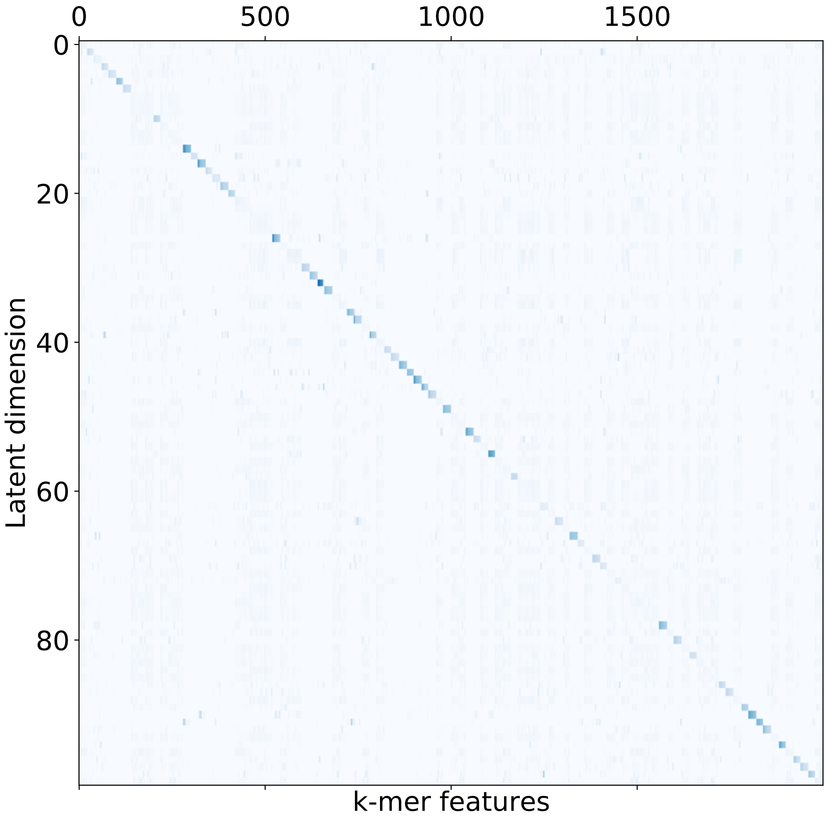


**Fig. S14:** Heatmap showing the top 20 k-mers learned by our model for each latent dimension in the A549 model

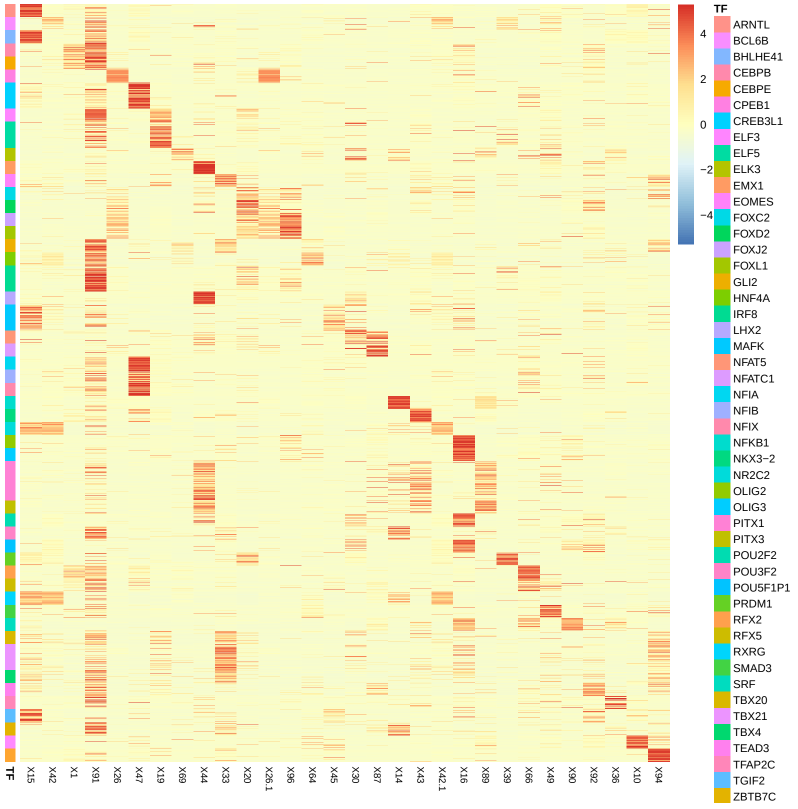


**Fig. S15:** Heatmap of the latent space obtained by our A549-trained model, upon doing inference on 11,600 SELEX probes from 51 TF experiments. Each row is the latent representation $\vec{z}$ of a HT-SELEX probe, with the rows being colored by the TF experiment that the probe comes from. There are 200 enriched probes per TF/HT-SELEX experiment.

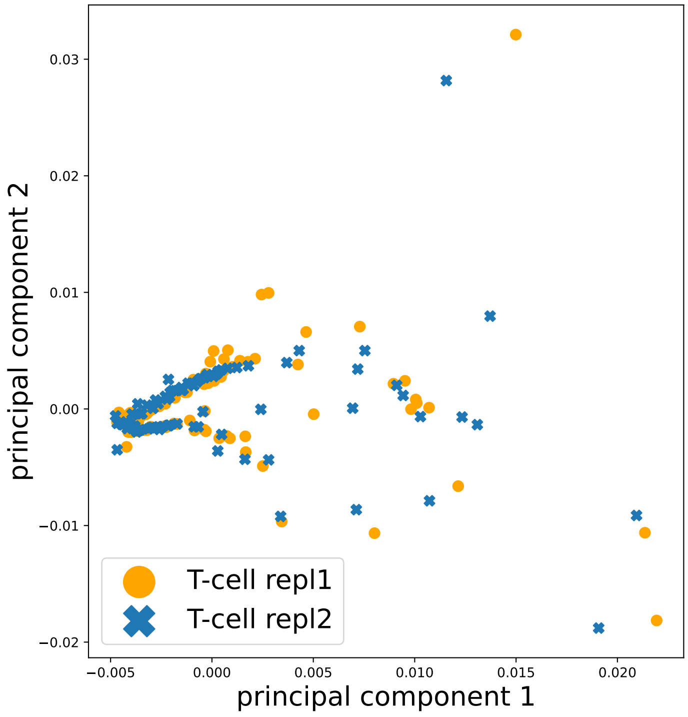


**Fig. S16:** PCA of the k-mer distributions in two isogenic replicates of T cells

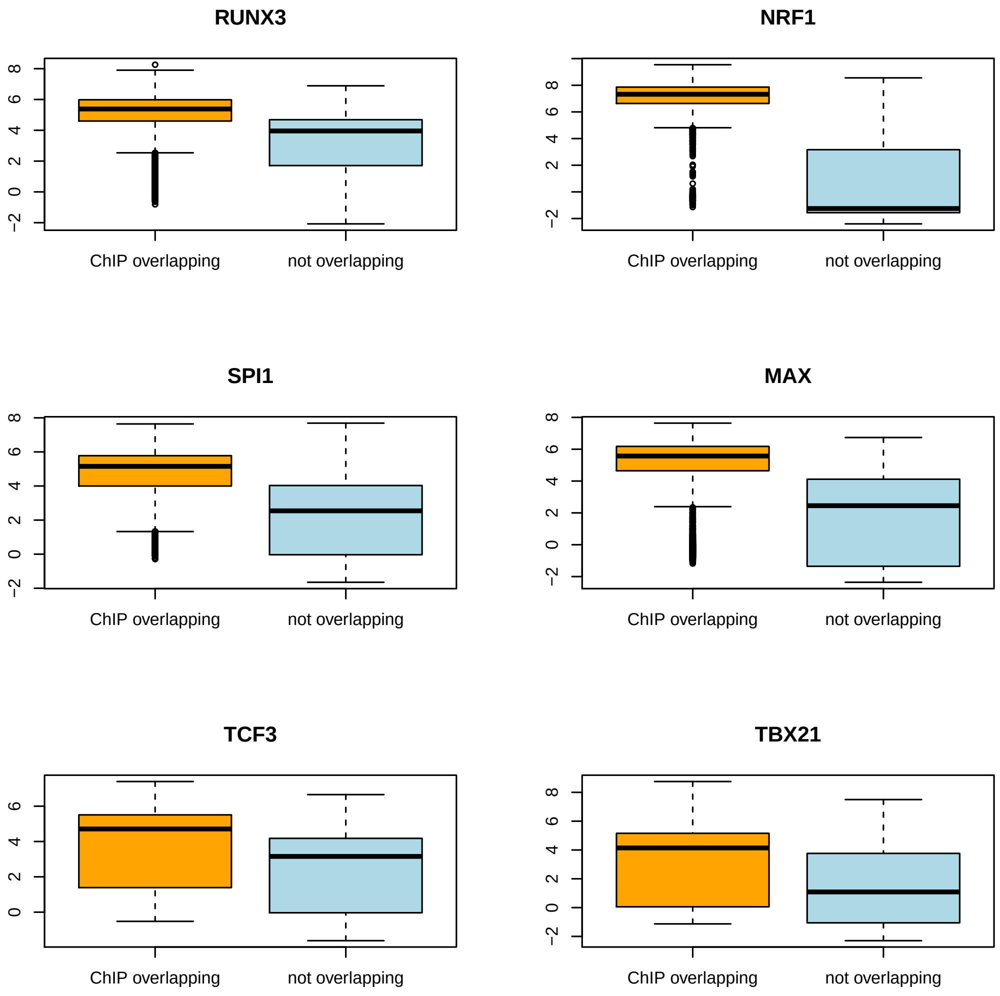


**Fig. S17:** Box-plots showing the distribution of topic scores for ATAC-seq peaks that overlap with ChIP-seq peaks and those that do not, for several TFs.

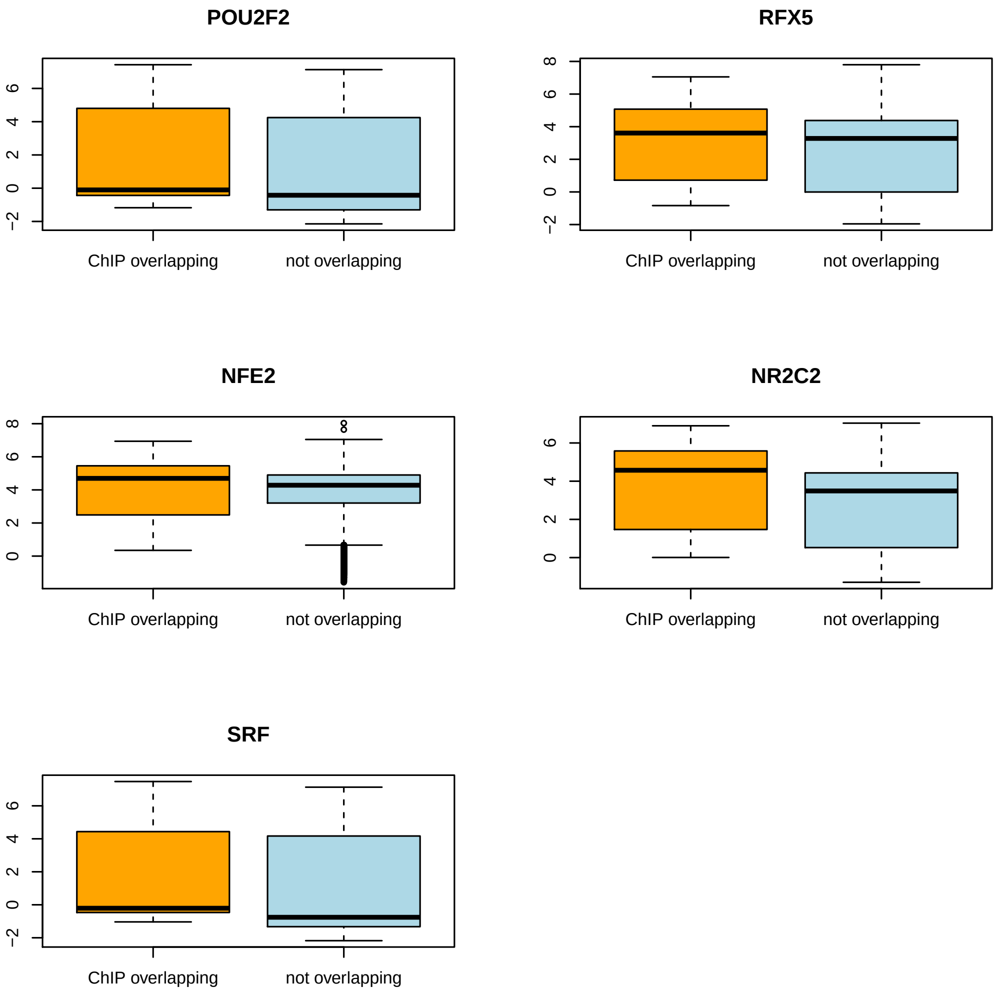


**Fig. S18:** Box-plots showing the distribution of topic scores for ATAC-seq peaks that overlap with ChIP-seq peaks and those that do not, for several TFs.
